# The G-protein-coupled estrogen receptor, a gene co-expressed with ERα in breast tumors, is regulated by estrogen-ERα signalling in ERα positive breast cancer cells

**DOI:** 10.1101/2022.06.14.496079

**Authors:** Uttariya Pal, Mohan Manjegowda, Neha Singh, Snigdha Saikia, Betty S. Philip, Deep Jyoti Kalita, Avdhesh Kumar Rai, Anupam Sarma, Vandana Raphael, Deepak Modi, Amal Chandra Kataki, Anil Mukund Limaye

## Abstract

**Purpose:** The purpose of this study was to assess the relationship between GPER, a seven transmembrane G-protein coupled estrogen receptor, and ERα in breast tumors, and to make inroads into the mechanistic basis and clinical significance.

**Methods:** TCGA-BRCA data was mined to examine the relationship between GPER and ERα expression. GPER mRNA, and protein expression were analyzed in ERα-positive or -negative breast tumors from two cohorts using immunohistochemistry, western blotting, or RT-qPCR. The Kaplan-Meier Plotter was employed for survival analysis. The influence of estrogen *in vivo* was studied by examining GPER expression levels in estrus or diestrus mouse mammary tissues, and the impact of 17β-estradiol (E2) administration in juvenile or adult mice. The effect of E2, or propylpyrazoletriol (PPT, an ERα agonist) stimulation on GPER expression was studied in MCF-7 and T47D cells, with or without tamoxifen or ERα knockdown. ERα-binding to the GPER locus was explored by analysing ChIP-seq data (ERP000380), in silico prediction of estrogen response elements, and chromatin immunoprecipitation assay.

**Results:** Clinical data revealed significant positive association between GPER and ERα expression in breast tumors. The median GPER expression in ERα-positive tumors was significantly higher than ERα-negative tumors. High GPER expression was significantly associated with longer overall survival of patients with ERα-positive tumors. *In vivo* experiments showed a positive effect of E2 on GPER expression. E2 induced GPER expression in MCF-7 and T47D cells; an effect mimicked by PPT. Tamoxifen or ERα-knockdown blocked the induction of GPER. Estrogen-mediated induction was associated with increased ERα occupancy in the upstream region of GPER.

**Conclusion:** GPER expression is positively associated with ERα in breast tumors, and a transcriptional target of the estrogen-ERα signalling axis. More in-depth studies are required to establish the significance of GPER-ERα co-expression, and their interplay in breast tumor development, progression, and treatment.

## INTRODUCTION

The estrogen receptor α (ERα) is a decisive variable that governs the mode of breast cancer treatment. It influences breast cancer prognosis and tumor phenotype. ERα expression is associated with indolence and favourable prognosis, whereas, its absence portends the opposite. Its expression in breast tumors also predicts response to endocrine therapy [1–3].

The genetic or molecular determinants of the aforementioned differences, and their relationship with ERα expression, is poorly understood. Arguably, the involvement of genes within the ERα co-expression network may hold the key [4]. On this supposition, Carmeci et al, employed differential screening of MCF-7 (ERα-positive), and MDA-MB-231 (ERα-negative) cDNA libraries to identify genes associated with ERα; only to find GPCR-Br, which was then an orphan G-protein-coupled receptor (GPCR), as an abundantly expressed marker in the former, but not in the latter. The study also demonstrated GPCR-Br and ERα co-expression in other breast cancer cell lines, and primary breast tumors [4]. Subsequent work from several laboratories showed that 17β-estradiol (E2) binds GPCR-Br [5, 6], to produce short-term non-genomic [5, 7], as well as long-term genomic effects [8, 9] of estrogen. Hence, it was re-christened as a G-protein-coupled estrogen receptor (GPER).

GPER signal transduction conforms to the typical GPCR signalling paradigm involving G_α_, and G_βγ_ subunits. The highlight of GPER signalling is the activation of EGFR/ERK/MAPK pathway via the HB-EGF ectodomain shedding [7], which not only induces proliferation [10], thereby mimicking EGF like effects, but also enables cross-talk with ERα via Ser118 phosphorylation [11] [12]. Tamoxifen is a GPER agonist [6]. Thus, GPER has gained traction in the field due to the following reasons: a) it serves as a non-canonical membrane estrogen receptor [5–7], b) it explains the EGF-like effects of estrogen [7], and c) it subserves tamoxifen resistance [13–15]. The acknowledgement of its clinical relevance is reflected in the plethora of publications addressing GPER expression in breast and other endocrine cancers, and its association with standard clinico-or histo-pathological markers [16, 17], patient survival [18, 19], endocrine resistance [15, 20], or metastasis [10].

The clinical import of ERα and GPER co-expression is of value, but warrants due attention in the face of the existing knowledge gaps. The co-expression, or association of GPER with ERα in breast tissue specimen was examined by several investigators. However, while some have reported positive association [16, 17, 21], others have reported negative [19, 22], or no association [15, 23, 24]. Hence, there is a need for independent assessment. Furthermore, the mechanistic basis, or significance of co-expression is unknown. Hormonal regulation of GPER, particularly by estrogen has been touched upon in several publications [13, 21]. However, the data are confusing, and do not address the role of hormone receptors, particulary the ERα.

The present study examined, and confirmed, positive association between ERα and GPER expression in breast tumor specimen from three independent cohorts. Higher expression in estrus, compared to diestrus mouse mammary tissues, and E2-driven, ERα-dependent modulation of GPER in human breast cancer cell lines revealed GPER as a transcriptional target of estrogen.

## MATERIALS AND METHODS

### Plasticwares and reagents

Cell culture plasticware was purchased from Eppendorf (Hamburg, Germany). Dulbecco’s modified eagle’s medium (DMEM) and Roswell Park Memorial Institute (RPMI) 1640 medium were purchased from HiMedia (Mumbai, India). Fetal bovine serum (FBS) was purchased from Invitrogen Corporation (Massachusetts, USA). Charcoal stripped fetal bovine serum (csFBS), Trypsin-EDTA, antibiotics, and Dulbecco’s phosphate-buffered saline (DPBS) were purchased from HiMedia (Mumbai, India). Anti-H3 antibody (Cat No. BB-AB0055) and Normal rabbit IgG antibody (Cat No. AB0001) was from BioBharati LifeScience (Kolkata, India). ERα (D8H8) antibody (Cat No. 8644S) and anti-rabbit secondary antibody (Cat. No. 7074S) were from Cell Signaling Technology (MA, USA). 17β-estradiol (E2, Cat No. E8875), 4-hydroxytamoxifen (TAM, Cat No. H7904), Propylpyrazoletriol (PPT, Cat No. H6036) and paraformaldehyde were purchased from Sigma (MO, USA). All other reagents, salts and buffers were purchased from Merck or SRL (Mumbai, India).

### TCGA-BRCA cohort

Publicly available breast cancer mRNA expression data from The Cancer Genome Atlas (TCGA), hereafter refered to as TCGA-BRCA data, were collected from the UCSC Xena Browser (xenabrowser.net). This data set contains genome-wide mRNA expression values in the form of log2(RPKM+1) on 114 normal breast tissues and 1097 primary breast tumors.

### Analysis of GPER expression in tumors from TCGA-BRCA cohort

As per the histogram shown in supplementary data 1b, ERα mRNA expression showed bimodal distribution in the primary breast tumors. The ERα mRNA expression data was modelled as a mixture of two Gaussian populations, which led to the estimation of mean ERα mRNA expression in ERα-low and ERα-high groups. Each tumor was classified as ERα-low or ERα-high if ERα mRNA expression was within two standard deviations (sd) of the estimated respective population means. The Shapiro-Wilk test revealed that ERα mRNA expression was not normally distributed in both the subgroups. Hence, Mann-Whitney U test was employed to compare median GPER mRNA expression in ERα-low and ERα-high tumors. Additionally, we considered the immunohistochemical data to classify primary breast tumors as ERα-positive or ER-negative, followed by comparison of the median GPER mRNA expression using the Mann-Whitney U test. We applied the Chi-squared test of significance to examine the association of GPER expression with ERα, PR status, molecular subtype, and stage.

### Survival analysis

Survival was analyzed using the Kaplan-Meier Plotter online (www.kmplot.com) [25]. The JetSet best probe “210640_s_at” for GPER was used for the analysis. Using the “auto select best cut-off” option, tumor samples were divided into GPER-high and GPER-low groups. The effect of GPER expression on overall survival (OS) of patients with ERα-positive or ERα-negative tumors, was graphically represented as Kaplan-Meier plots.

### Breast tumor samples

GPER protein expression was examined in a sample of 65 archival paraffin embedded breast tumors, from the Department of Pathology, North Eastern Indira Gandhi Regional Institute of Health and Medical Sciences (NEIGRIHMS), Shillong, India. These samples were from a cohort of breast cancer patients registered in NEIGRIHMS from 2007 to 2016. The mean age of the cohort was 47 years (range: 26-86 years). Additionally, breast tumors were also collected from patients registered at Dr. Bhubaneswar Borooah Cancer Institute (BBCI), Guwahati, India from December 2018 to October 2019, after obtaining patients’ informed consent, which were used for analysis of GPER mRNA and protein expression. The mean age of the cohort was 47.3 years (range: 26-82 years). ER, PR, and HER-2 expression data were available for the samples. The studies were approved by ethics committees of both centers.

### Animals and treatments

Cyclicity of 8-10 weeks old female C57BL/6J mice was monitored by examination of vaginal swabs stained with Giemsa (Cat No. SO11, Himedia, Mumbai, India). The animals in estrus (n = 11) or diestrus stages (n = 8) of the reproductive cycle were euthanized before collecting the mammary tissues for analysing Gper mRNA and protein using RT-qPCR and immunofluorescence.

To study the effects of estrogen, 5 weeks and 8 weeks old C57BL/6J mice were administered with E2 (Cat No. E-2758, Sigma, MO, USA), which was dissolved in ethanol and diluted in corn oil (Cat No. C8267, Sigma, MO, USA) at a dose of 1µg/day/mouse subcutaneously for 5 consecutive days. Controls were administered equal amounts of ethanol in corn oil. On 6th day afternoon, the animals were euthanized and mammary tissues were collected. The study was replicated thrice with 3 animals in each group. The study was approved by institutional animal ethics committee (IAEC Project No. 1/16).

### Immunohistochemistry

Sections of 4 µm thickness were cut from the paraffin-embedded tissues and mounted on silane-coated microscopic slides. They were deparaffinised with xylene after an overnight incubation in a hot air oven set at 37°C. The sections were rehydrated with decreasing concentrations of ethanol in water, before microwaving for 10 min at 450 W, followed by 15 min at 600 W in citrate buffer (10 mM, pH 6) for antigen retrieval. The slides were allowed to cool and then rinsed once with PBS. Immunohistochemical staining was done using Super Sensitive™ IHC detection system from BioGenex Laboratories (CA, USA) as per the manufacturer’s instructions. Briefly, endogenous peroxidase activity was quenched by peroxide block for 10 min followed by three PBS washes of 5 min each. Sections were then probed with primary antibody [26] for 1 h at a dilution of 1:50. Sections were then incubated with super enhancer (20 min), polymer-HRP (30 min) and DAB substrate (10 min). After each step, the excess reagent was removed by three PBS washes of 5 min each. DAB staining was stopped by rinsing the slides in running tap-water. Sections were counter-stained with Harris hematoxylin and dehydrated by incubating in increasing concentrations of ethanol followed by xylene. Slides were air dried and mounted using DPX mountant, and observed under the microscope.

Modified semi-quantitative scoring of IHC for GPER was done only for the cytoplasmic compartment. The staining intensity was classified as negative (-), very weak (1+), weak (2+) and strongly positive (3+). The H-score was calculated by multiplying the intensity score with the percentage of positive cells with a score ranging from 0 to 300. Thus, the H-score took into consideration both the staining intensity and the percentage of cells. In every batch of slides, normal tissue within the biopsy was taken as internal control and slide with no primary antibody was taken as negative control. In the present study, though occasional tumor samples showed membrane and nuclear positivity, GPER staining was predominantly cytoplasmic. The H-score of the normal breast tissue was less than 100 in 29/35 cases and mostly ranged from 80-90. Samples with H-score > 40 were considered as GPER-positive.

Specificity of staining was confirmed by a peptide blocking experiment involving 4 slides of serial sections. They were incubated with anti-GPER antibody (+ve control), anti-GPER antibody pre-incubated with the peptide (peptide blocked antibody), anti-GPER antibody with peptide resuspension buffer (resuspension buffer control), or no primary antibody (-ve control) for 1 h before proceeding with secondary antibody incubation and chemiluminescence detection. As shown in supplementary data 2, absence of staining in samples treated with the antibody pre-incubated with the peptide indicated the specificity of staining.

### Immunofluorescence

Mammary tissues were harvested and washed in PBS. They were stretched on glass slides, air dried and fixed in 4% paraformaldehyde. Paraffin embedded mouse mammary tissue sections of 5 µm thickness were deparaffinized and hydrated in grades of methanol. Epitopes were exposed in antigen retrieval buffer (Tris-EDTA, pH-9.0) for 35 minutes at 95⁰C. Slides were incubated in Gper antibody (Cat No. sc-48524-R, Santa Cruz Biotechnology, Texas, USA) using 1:100 dilution at 4⁰C, overnight. The slides were then washed in 1X PBS for 30 minutes followed by incubation with biotinylated secondary antibody for 40 min at room temperature. Signal was amplified by streptavidin-conjugated FITC (Cat No. FTC9, Bangalore Genei, Bangalore, India, Dilution 1:100) at RT for an hour. Nuclei were counterstained with DAPI (Sigma-Aldrich, Missouri, USA) and mounted in VECTASHIELD Antifade media (Cat No. H-1200-10, Vector Laboratories, CA, USA). Slide edges were sealed with nail polish, and imaged with an oil immersion Plan-Apochromat 63X/ 1.4 NA objective using a confocal laser scanning microscope (Zeiss, Jena, Germany) at IIT Bombay.

### Cell culture

MCF-7 and T47D breast cancer cell lines were procured from the National Centre for Cell Science, Pune, India. Phenol red-containing DMEM or RPMI-1640, supplemented with 10% heat-inactivated FBS, 100 μg/ml streptomycin and 100 units/ml penicillin (M1 medium) was used for routine culture of MCF-7, or T47D cells, respectively. Cells were cultured under humidified conditions of 5% CO_2_ at 37°C. At 70–80% confluency, cells were trypsinized and subcultured, or seeded for experiments.

### Treatment of cell lines

2 × 105 MCF-7 or T47D cells were seeded in 6-well plates, in M1 medium. After 48 h of incubation, the monolayer was washed with DPBS, and fed with phenol red-free DMEM (for MCF-7) or RPMI-1640 (for T47D), supplemented with 10% heat-inactivated csFBS, 100 units/ml penicillin, and 100 µg/ml streptomycin (M2 medium). After 24 h, the cells were washed with DPBS, and treated in M2 medium, with indicated concentrations of E2, PPT, or tamoxifen, alone or in combinations, for indicated period of time. Control cells were treated with 0.1% ethanol in M2 medium. The treatment medium was replenished every 48 h.

### siRNA transfection

2 × 105 MCF-7 or T47D cells were seeded in 6-well plates and incubated for 24 h in M1 medium. Cells were transfected with ERα-specific (Cat No. 4392420, Thermo Scientific, PA, USA) or scrambled siRNA (Cat No. AM4611, Thermo Scientific, PA, USA) using Lipofectamine RNAiMAX (Cat No. 13778075, Thermo Scientific, PA, USA) for 48 h according to the manufacturer’s instructions.

### RNA extraction, cDNA synthesis and RT-qPCR

The mRNA expression of GPER variants in breast tumor samples, and treated or untreated MCF-7 or T47D cells was analyzed by RT-qPCR. Total RNA was extracted and reverse-transcribed as described earlier [27] and cDNA equivalent to 20 ng of total RNA was amplified with gene specific primers (table 1). RT-qPCR reactions were carried out in AriaMX (Agilent, CA, USA). Reactions were set up in 1X PowerUP SYBR Green master mix (Cat No. A25743, Thermo Scientific, PA, USA). ROX dye served as passive reference. RPL35a served as an internal control.

**Table 1.**
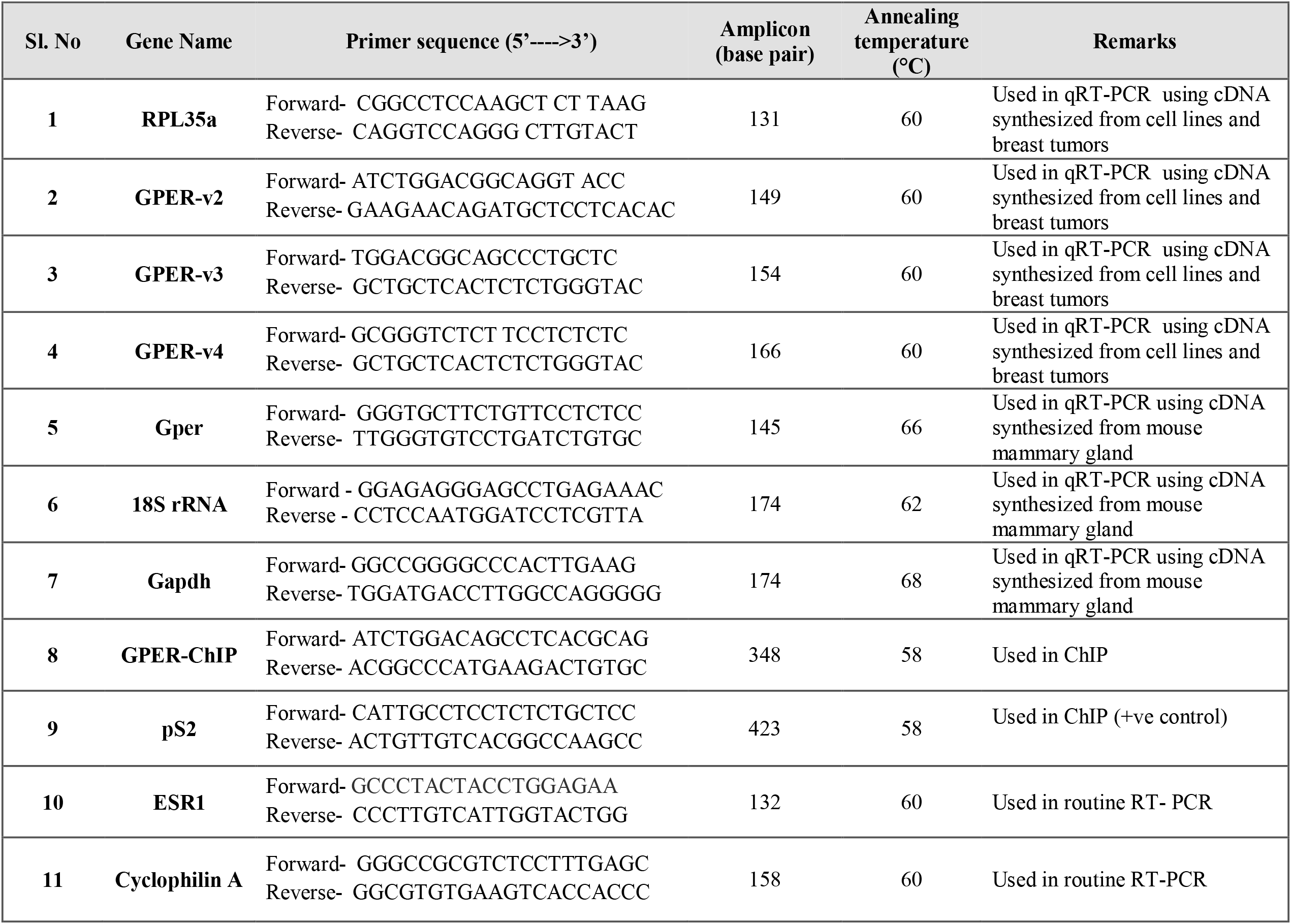
List of primers used for routine RT-PCR, RT-qPCR, and ChIP.

For the analysis of Gper mRNA expression in mouse mammary tissues, total RNA was extracted with TRIzol (Invitrogen, Massachusetts, USA), as per manufacturer’s instructions, precipitated in isopropanol, and washed with 70% alcohol, dried, and dissolved in 20 µl of DEPC water. Total RNA was treated with DNase I (Bangalore Genei, Bangalore, India) as described previously [28] and cleaned using RNeasy mini kit (Qiagen, Hilden, Germany). RNA was quantified by nano-spectrophotometer (Eppendorf, Hamburg, Germany). 1 µg of RNA was reverse transcribed using High-capacity cDNA Reverse Transcription Kit (Applied Biosystems, Massachusetts, USA) as mentioned previously [29]. The cDNA was diluted 10 times, and RT-qPCR reaction was run using 2 µl of diluted cDNA in duplicates against Gper primer (table 1). RT-qPCR was performed using iQ SYBR green super mix in the CFX-96 thermal cycler (Bio-Rad, CA, USA). Amplification was carried out for 35 cycles where each cycle involved initial denaturation 95°C for 30 sec, primer annealing for 30 sec, and extension at 72°C for 30 sec. The fluorescence emitted was collected for 30 sec in the extension step of each cycle. The homogeneity of the PCR amplicons was verified using the melt curve method. All PCR amplifications were carried out in duplicate, for all biological replicates. Gene expression was normalized to 18S rRNA and estimated using Pfaffl method [30].

### Western blotting

Total protein was isolated, quantified and subjected to western blotting with peptide affinity purified antibody to analyze GPER protein expression in breast tumor samples and cell lines as described previously [26]. Histone (H3) served as the internal control.

### ChIP-seq analysis

Sequence Read Archive (SRA) was searched for ERα-related ChIP-seq studies in MCF-7 cells. Subset of files from the project ERP000380 was selected. The raw data quality was assessed in galaxy web based platform [31]. The quality of all the input read files were assessed by FASTQC [32] with the default settings. The quality scores were converted to Sanger quality type by FASTQ Groomer [33]. Reads were mapped to reference human genome (hg19) using “Map with Bowtie for Illumina” tool [34]. Reads mapping to multiple locations were discarded and unmapped reads in the output SAM file were filtered using “Filter SAM or BAM, output SAM or BAM” tool [35]. The genomic regions enriched by reads were identified by MACS (Model-based analysis of ChIP-Seq) tool [36]. The peaks were visualized in UCSC genome browser [37] after converting Wig files to bigWig files using “Wig/BedGraph-to-bigWig” tool.

### Chromatin immunoprecipitation (ChIP) assay

The ChIP protocol used in this study was as described earlier [29]. Briefly, cells were fixed with 0.75% (v/v) formaldehyde at room temperature for 10 mins followed by addition of 125 mM glycine to inhibit the cross-linking reaction. Cells were washed and lysed with 1 ml of ChIP lysis buffer. The lysates were sonicated, to shear the DNA at an amplitude of 30% for 35 cycles, each cycle with a 10-sec pulse on and 25-sec pulse off. Sonicated samples were centrifuged at 14000 rpm for 10 mins at 4°, and the supernatant containing the sheared chromatin was pre-cleared with Protein G plus-Agarose beads (Cat No. IP04-1.5ML, Merck Millipore, Burlington, USA), which were pre-coated with bovine serum albumin (BSA, Cat No. PG-2330, Puregene, CA, USA) and herring sperm DNA (Cat No. D7290, Sigma, Missouri, USA). 5% of the pre-cleared chromatin samples were kept aside as input. The remaining portion of the chromatin was incubated with normal rabbit IgG antibody or ERα antibody for 4 h. Protein-antibody complexes were immunoprecipitated by incubating with 20 µl of pre-coated Protein G plus-Agarose beads followed by centrifugation. After washing the beads extensively, the DNA was eluted in elution buffer, column purified and used as a template in PCR reactions with primers to amplify a specific region of GPER (test) or TFF1 (pS2) (positive control, [29]) locus.

### Statistical analysis

Two-group, non-normal data were analyzed by Mann-Whitney U test to examine significant differences in median. Else, Welch two-samples t-tests were used. Chi-squared tests, or Fisher’s exact tests were used as indicated to test for significance in the association between GPER expression with histo-or clinicopathological variables. Multiple group data were analysed by one-way ANOVA, followed by Tukey’s HSD post-hoc test. Statistical tests were carried out in R, considering 5% level of significance (p < 0.05).

## RESULTS

The inconsistency in the literature on the association between ERα and GPER, motivated us to examine their correlation in primary breast tumors (n = 1097) from the TCGA-BRCA cohort. There was a significant correlation between the two variables (Pearson’s r = 0.266, p < 0.0001, supplementary data 1a). However, the existence of two subgroups of breast tumors, namely ERα-low and ERα-high, was apparent as illustrated in supplementary data 1b. We determined the mean ERα mRNA expression in these subgroups by mixture modelling. ERα-low (n = 286), and ERα-high (n = 790) tumors were considered as those that had ERα mRNA expression levels within two standard deviations on either side of their respective means (Fig 1a). There was a significant difference in the median GPER mRNA levels in the subgroups; the ERα-high tumors expressing higher median levels (Fig 1b). Median GPER mRNA expression was also significantly different in immunohistochemically classified ERα-positive, and ERα-negative breast tumors of the TCGA-BRCA cohort (Fig 1c). Table 2 shows that GPER-high tumors were more frequent in ERα-, or PR-positive breast tumors, as compared to the negative ones. A Chi-squared test revealed that GPER mRNA expression was significantly associated with ER, or PR positivity (p < 0.0001).

**Fig 1:**
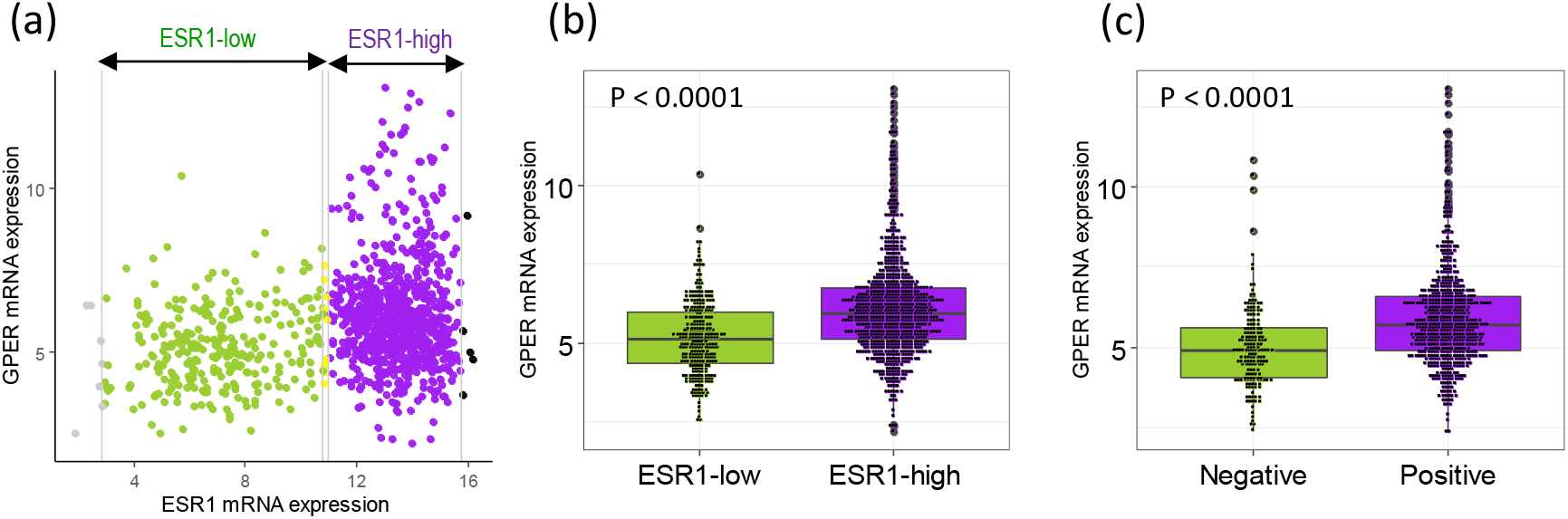
GPER expression in TCGA-BRCA breast tumors stratified according to ERα expression. A scatter-plot of GPER and ERα mRNA levels in primary breast tumors (n = 1097). Note the bimodality of ERα expression, which has been depicted as a histogram in supplementary data 1b. The ERα expression was modelled as a mixture of two Gaussian populations. The green and blue colored clusters represent ERα-low and ERα-high tumors, respectively, which were grouped on the basis of expression level within two standard deviations on either side of the respective means. (b) A boxplot showing the distribution of GPER mRNA expression in ERα-low (n = 286) and ERα-high (n = 790) breast tumors identified in (a). (c) A boxplot showing the distribution of GPER mRNA expression levels in ERα-negative (Negative, n = 179) and ERα-positive (Positive, n = 601) breast tumors on the basis of immunohistochemical assessment. ESR1 symbol for gene encoding human ERα. mRNA expression levels are in terms of log_2_>(RPKM+1) values. Data in (b) and (c) were analysed by Mann-Whitney U test (***p < 0.0001)

**Table 2.** Association of the GPER expression with the histopathological parameters (TCGA-BRCA cohort).

|  | **GPER-high** | **GPER-low** | **p** |
| --- | --- | --- | --- |
| **Age** |  |  |  |
| Mean ± SD | 57.6 ± 14.0 | 58.3 ± 12.7 | T: 0.4875 |
| Median | 58 | 58 |  |
| Range | 26-90 | 31-90 |  |
| **ERα** |  |  |  |
| ERα-positive | 319 (57.48) | 236 (42.52) | < 0.0001 |
| ERα-negative | 39 (24.22) | 122 (75.78) | |
| **PR** |  |  |  |
| PR-positive | 283 (58.23) | 203 (41.77) | < 0.0001 |
| PR-negative | 75 (32.61) | 155 (67.39) |  |
| **Molecular type** |  |  |  |
| Normal-like | 9 (50.00) | 9 (50.00) | <0.0001 |
| Luminal A | 233 (66.38) | 118 (33.62) |  |
| Luminal B | 74 (44.58) | 92 (54.42) |  |
| Basal-like | 32 (26.02) | 91 (73.98) |  |
| HER2-enriched | 10 (17.24) | 48 (82.76) |  |
| **Tumor Stage** |  |  |  |
| Stage I | 68 (56.67) | 52 (43.33) | 0.2075 |
| Stage II | 196 (47.80) | 214 (52.20) |  |
| Stage III | 85 (52.47) | 77 (47.53) |  |
| Stage IV | 4 (28.57) | 10 (71.43) |  |
| Stage X | 5 (50.00) | 5 (50.00) |  |

Number within the braces indicates % of GPER-high or -low in various categories. The p-values (p) are from Chi-squared test, T: student's t-test. In all the tests $p < 0.05$ was considered as significant.

We also employed immunohistochemistry to examine GPER protein expression in archival paraffin embedded breast tumor specimens from NEIGRIHMS cohort using a polyclonal antibody raised against the N-terminal peptide of GPER [26]. The specificity of the antibody was demonstrated by a peptide blocking experiment (supplementary data 2). Representative images showing varied intensities of GPER staining in tumors with different expression levels of ER, PR and HER2 are provided as a supplement (supplementary data 3). H-score of 40 was used as a cut-off to segregate the GPER-positive (H > 40) from the GPER-negative tumors (H<=40). 44 out of 65 (68%) of the tumors were GPER-positive, whereas the remaining (32%) were GPER-negative (table 3). A Chi-squared test showed significant association between GPER expression and ERα. The tumors were also classified based on TNM stage, Bloom Richardson grade, lymphovascular invasion, tumor type and size, margin type, lymph node status, and molecular subtype. 21 out of 44 (47%) GPER-positive tumors, and 4 out of 21 (19%) GPER-negative tumors were positive for lymphovascular invasion, suggesting a positive association between GPER expression and lymphovascular invasion (table 4, p = 0.031). None of the other clinicopathological parameters was significantly associated with GPER expression in this study (table 4).

**Table 3.** Association of GPER expression with immunohistochemical markers in tumor samples of the NEIGRIHMS cohort.

|  | GPER-positive<br>(44/65) | GPER-negative<br>(21/65) | p |
| --- | --- | --- | --- |
| **ERα** |  |  |  |
| Positive | 33 | 10 | 0.048 |
| Negative | 11 | 11 |  |
| **PR** |  |  |  |
| Positive | 32 | 9 | 0.028 |
| Negative | 12 | 12 |  |

The p values (p) are from Chi-squared test and $p < 0.05$ was considered as significant.

**Table 4.** GPER expression in tumor samples of the NEIGRIHMS cohort and its association with clinicopathological parameters.

|  | GPER-positive<br>(44/65) | GPER-negative<br>(21/65) | p |
| --- | --- | --- | --- |
| **TNM stage** |  |  |  |
| Early (Stage 1 & 2) | 21 | 9 | F: 0.79 |
| Advanced (stage 3 & 4) | 23 | 12 | C: 0.92 |
| **Bloom Richardson Grade** |  |  |  |
| 1 | 11 | 4 | F: 0.838 |
| 2 | 18 | 8 |  |
| 3 | 15 | 9 |  |
| **Lymphovascular invasion** |  |  |  |
| Positive | 21 | 4 | F: 0.031 |
| Negative | 23 | 17 |  |
| **Age** |  |  |  |
| Mean ± SD | 47 ± 11.95 | 48.9 ± 11.17 | T: 0.55 |
| Median | 46.5 | 48 | M: 0.77 |
| Range | 26 to 86 yrs | 35 to 86 yrs |  |
| **Tumor type** |  |  |  |
| IDC | 39 | 20 | F: 0.65 |
| Others | 5 | 1 |  |
| **Margin type** |  |  |  |
| Invasive | 39 | 19 | F: 1.00 |
| Pushing | 5 | 2 |  |
| **Tumor size** |  |  |  |
| Mean ± SD | 4.020 ± 1.896 | 4.476 ± 1.735 | T: 0.355 |
| Median | 3.75 | 4 | M: 0.246 |
| Range | 1.5 to 11 cm | 1.5 to 6 cm |  |
| **Lymph node status** |  |  |  |
| Involved | 23 | 12 | F: 0.79 |
| Uninvolved | 21 | 9 |  |
| **Molecular type** |  |  |  |
| Luminal A | 20 | 7 | F: 0.115 |
| Luminal B | 13 | 3 |  |
| Triple negative | 2 | 4 |  |
| HER2/neu type | 9 | 7 |  |

The p values are from C: Chi-squared test, T: student's t-test, F: Fisher's exact test, M: Mann-Whitney U test. In all the tests $p < 0.05$ was considered as significant.

We applied western blotting technique to study GPER protein levels in immunohistochemically confirmed ERα-positive or ERα-negative breast tumors received from BBCI cohort. A representative western blot in Fig 2a shows higher GPER protein levels in ERα-positive breast tumors. GPER protein is encoded by three annotated transcript variants, referred to as GPER-v2 (NM_001505.2), -v3 (NM_001039966.1), and -v4 (NM_001098201.1). These transcripts have the same coding sequence and a 3’-UTR, but possess different 5’-UTRs. A previous work from our laboratory showed that the aforementioned variants are expressed in breast cancer cell lines; MCF-7 cells expressing elevated levels compared to MDA-MB-231 [27]. Recently we reported another variant of GPER, referred to as GPER-v5 [38]. We examined the expression of these transcript variants in immunohistochemically confirmed ERα-positive (n = 25) and ERα-negative (n = 25) breast tumors of the BBCI cohort. As shown in Fig 2b, GPER-v2, -v3 and -v4 were significantly elevated in ERα-positive breast tumors compared to the negative ones. We were unable to detect the expression of GPER-v5 in these samples.

**Fig 2:**
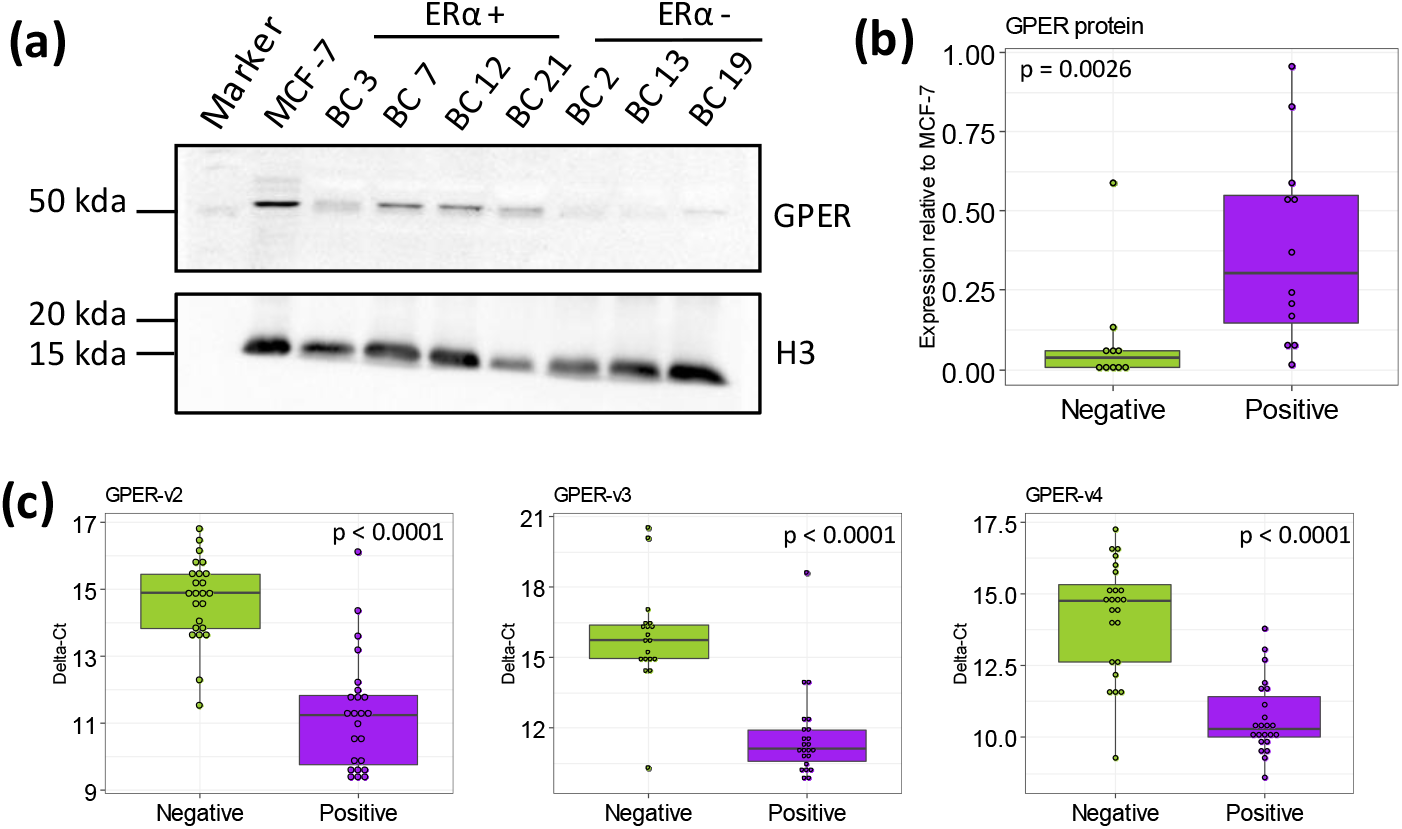
GPER protein and mRNA expression in ERα-negative and ERα-positive breast tumors from the BBCI cohort. (a) A representative western blot showing GPER protein levels in ERα-positive (n = 4) and ERα-negative (n = 3) breast tumors. The blots were probed with a peptide affinity purified antibody against GPER [26]. (b) Quantitative representation of GPER protein expression in ERα-negative (n = 10) and ERα-positive (n = 12) breast tumors. Chemiluminescence signals were processed and quantified using ImageJ [53]. For each sample, the background-subtracted integrated band intensity for GPER was normalized against that obtained for Histone (H3). Normalized GPER protein expression in tumor samples was expressed relative to that in MCF-7 cells, which was set to 1. Data were analyzed by Mann-Whitney U test test (*p < 0.05). The chemiluminescence data for all the samples analysed are compiled as supplementary data 4. (c) GPER mRNA variant expression levels in ERα-negative and ERα-positive breast tumors. Total RNA isolated from ERα-positive (n = 25) and ERα-negative (n = 25) breast tumors were reverse transcribed, and the resultant cDNAs were subjected to RT-qPCR using variant specific primers. RPL35a was used as an internal control. For each sample the average Ct value for a given mRNA variant (Ctvariant), and RPL35a (CtRPL35a) obtained from duplicate reactions were determined. The difference Ctvariant -CtRPL35a, was considered as a measure of the normalized variant expression. Samples with no Ct values in duplicate technical replicates were excluded, GPER-v2 (ERα-positive, n = 24; ERα-negative, n = 24), GPER-v3 (ERα-positive, n = 23; ERα-negative, n = 19) and GPER-v3 (ERα-positive, n = 23; ERα-negative, n = 24). Box plots show the normalized GPER variant mRNA expression in the BBCI cohort. Data were analyzed by Mann-Whitney U test. Note that the difference is significantly lower in ERα-positive tumors, indicating higher expression.

Previous studies have shown that primary breast tumors within the TCGA-BRCA cohort exhibit significantly lower levels of GPER mRNA expression than normal breast tissues [27]. Higher GPER expression in breast tumors were shown to have a positive effect on patient survival [27]. The aforementioned results motivated us to examine the effect of GPER-ERα co-expression on patient survival. As shown in Fig 3a, high GPER expression in ERα-positive tumors was significantly associated with prolonged OS. In contrast high GPER expression in ERα-negative tumors evidently had an opposite effect (Fig 3b).

**Fig 3:**
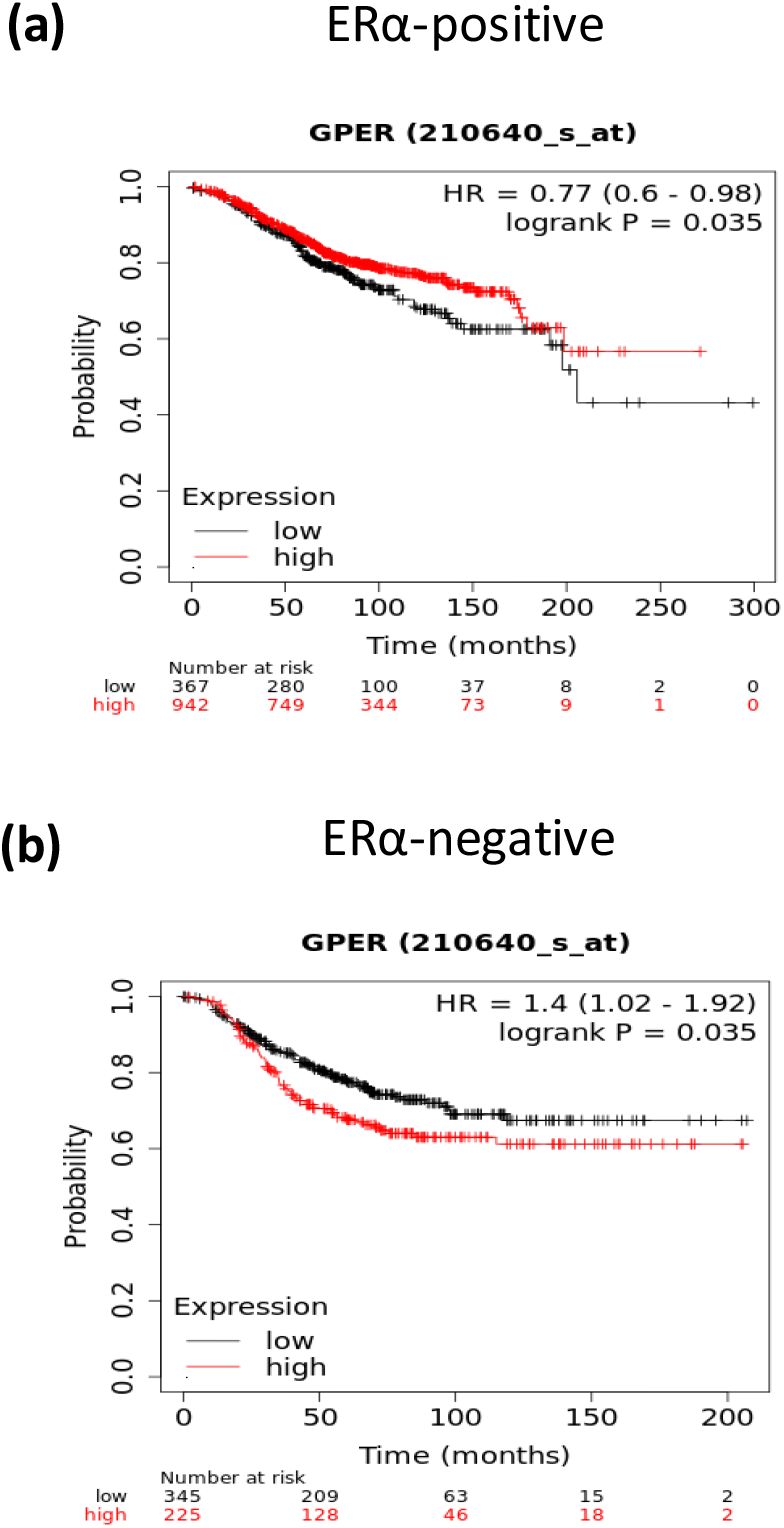
The influence of GPER expression on overall survival depends on ERα status of breast tumors. Relationship between GPER expression and survival of breast cancer patients. Survival analyses were performed using Kaplan–Meier Plotter online tool [25]. The JetSet best probe “210640_at” was considered for GPER expression. The breast tumors were classified as GPER-high or -low based on “auto select best cut-off” option. OS of patients stratified on the basis of high, or low GPER expression in patients with ERα-positive (a) or ERα-negative (b) breast tumors separately.

We studied Gper protein and mRNA expression in the mouse mammary tissues at estrus and diestrus stages. Immunostaning for Gper was detected in both alveolar and ductal epithelium. At the diestrus stage when the glands are smaller, Gper was weak and restricted mainly to the apical region of ductal epithelial cells. At the estrus stage, when the epithelial cells were multi-layered, glands grow in size, abundance of Gper protein per cell was high and strong in the epithelium. The expression of Gper in estrus stage was mainly cytoplasmic and not polarized like that at the diestrus stage (Fig 4a). The staining in the epithelium is specific as no reactivity was noted in these cells when primary antibody was replaced by normal rabbit serum. The staining in the stroma is non-specific as it was detected also in the negative controls. (Fig 4a). RT-qPCR analysis results also corroborate with the immunostaining results and reveal that expression of Gper mRNA was 2 fold higher in estrus stage when compared to diestrus (Fig 4b).

**Fig 4:**
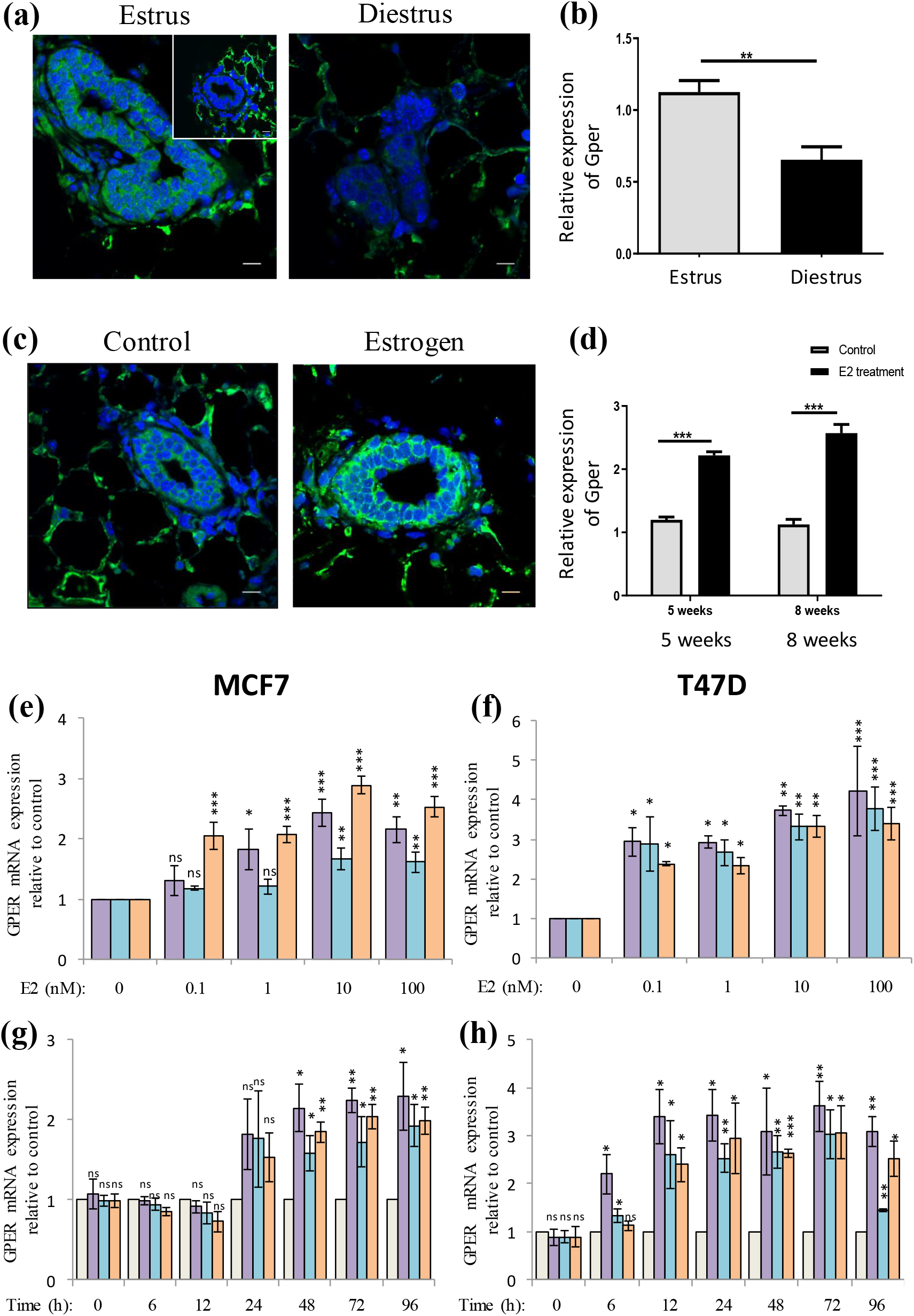
Estrogen positively affects GPER expression. (a, b) Immunofluorescence (a) and RT-qPCR study of Gper protein and mRNA expression respectively in mouse mammary tissues during estrus and diestrus phases of the reproductive cycle. Immunostaining for Gper (Green channel), and DAPI stained nuclei (blue channel). Bar is 50 microns. No antibody control is in inset. Mammary tissue was collected from adult cycling mice in estrus (n = 11) & diestrus (n = 8) stages and processed for RT-qPCR. Graph is plotted as mean ± sd. Y-axis shows normalized relative expression. p < 0.05 was taken as statistically significant. **p < 0.01. (c,d) Immunofluorescence (c), and RT-qPCR (d) study of the effect of 17β-estradiol treatment on Gper protein, and mRNA, respectively, in juvenile (5 weeks), or adult (8 weeks) old mice. Juvenile female mice were treated with estrogen for five days and mammary tissue was collected. Immunostaining for Gper (Green channel), and DAPI stained nuclei (blue channel) was performed. Bar is 50 microns. (d) Juvenile (5 weeks) and adult female (8 weeks) mice were treated with estrogen for five days and breast tissue was subjected to RNA extraction and RT-qPCR. Graph is plotted as mean ± sd. Y-axis shows normalized relative expression. p < 0.05 was taken as statistically significant. ***p < 0.001. (e, f) A dose-response study of the effect of E2 treatment on GPER mRNA expression in MCF-7 (e), and T47D cells (f). Cells were treated with indicated concentrations of E2 for 72 h. Expression of GPER mRNA variants relative to RPL35a was analyzed using RT-qPCR. Expression levels in vehicle controls (ethanol treated cells) were set to 1 and those in E2 treated groups were expressed relative to control. Bars represent mean ± sd (n = 3). Data for each variant were subjected to ANOVA with Tukey’s HSD. *p < 0.05, **p < 0.01, ***p < 0.001. (g, h) A time-course study of the effect of E2 on MCF-7 (g) or T47D (h) cells. Cells were treated with 10 nM E2 or vehicle for indicated durations. For each sample the expression of GPER mRNA variants relative to RPL35a was analyzed using RT-qPCR. For each time-point the GPER variant expression was expressed relative to vehicle control, which was set to 1. Bars represent mean ± sd (n = 3). For each time-point data were analyzed by Welch two-sample t-test. *p < 0.05, **p < 0.01, ***p < 0.001. Colored bars correspond to GPER-v2 (purple), GPER-v3 (light blue), and GPER-v4 (orange) variant mRNAs. The grey bars (g,h) represent control cells, and are included to show that each time point had its ethanol treated control.

To test if estrogen induces expression of Gper, we isolated mammary glands from juvenile (5 weeks) and adult mice (8 weeks) treated with 17β-estradiol. Gper expression was increased with E2 treatment and protein was cytoplasmic (Fig 4c). As compared to untreated controls Gper mRNA levels increased significantly in both juvenile and adult mice (Fig 4d). This increase was statistically significant (p-value <0.001).

The aforementioned *in vivo* results motivated a mechanistic study of the estrogen regulation of GPER in human breast cancer cell lines. Dose-response (Fig 4, e,f) and time-course experiments (Fig 4, g,h) confirmed estrogenic induction of GPER mRNA variants in MCF-7 and T47D cells. PPT, a synthetic non-steroidal ERα agonist, also induced GPER mRNA variants in both the cell lines (Fig 5, a,b). Tamoxifen blocked estrogen-or PPT-mediated induction (Fig 5, a,b); a strong indication of ERα involvement, which was confirmed by ERα knockdown experiments in both the cell lines (Fig 5, c,d). Liganded ERα modulates gene expression by engaging with estrogen response elements in target gene promoters. We hypothesized that GPER is a direct transcriptional target of liganded ERα. To test this hypothesis, we analyzed ChIP-seq data obtained from MCF-7 cells treated by E2 for 24 h. Several ERα enriched regions in the GPER locus were apparent (Fig 5e). Furthermore, both MATINSPECTOR, and JASPAR tools predicted an ERE in the region corresponding to the 5’-most peak revealed in the ChIP-seq data. A ChIP assay employing ERα specific antibody, and primers designed to amplify this region, confirmed the enhanced ERα occupancy in MCF-7 cells exposed to 10 nM E2 for 24 h (Fig 5f).

**Fig 5:**
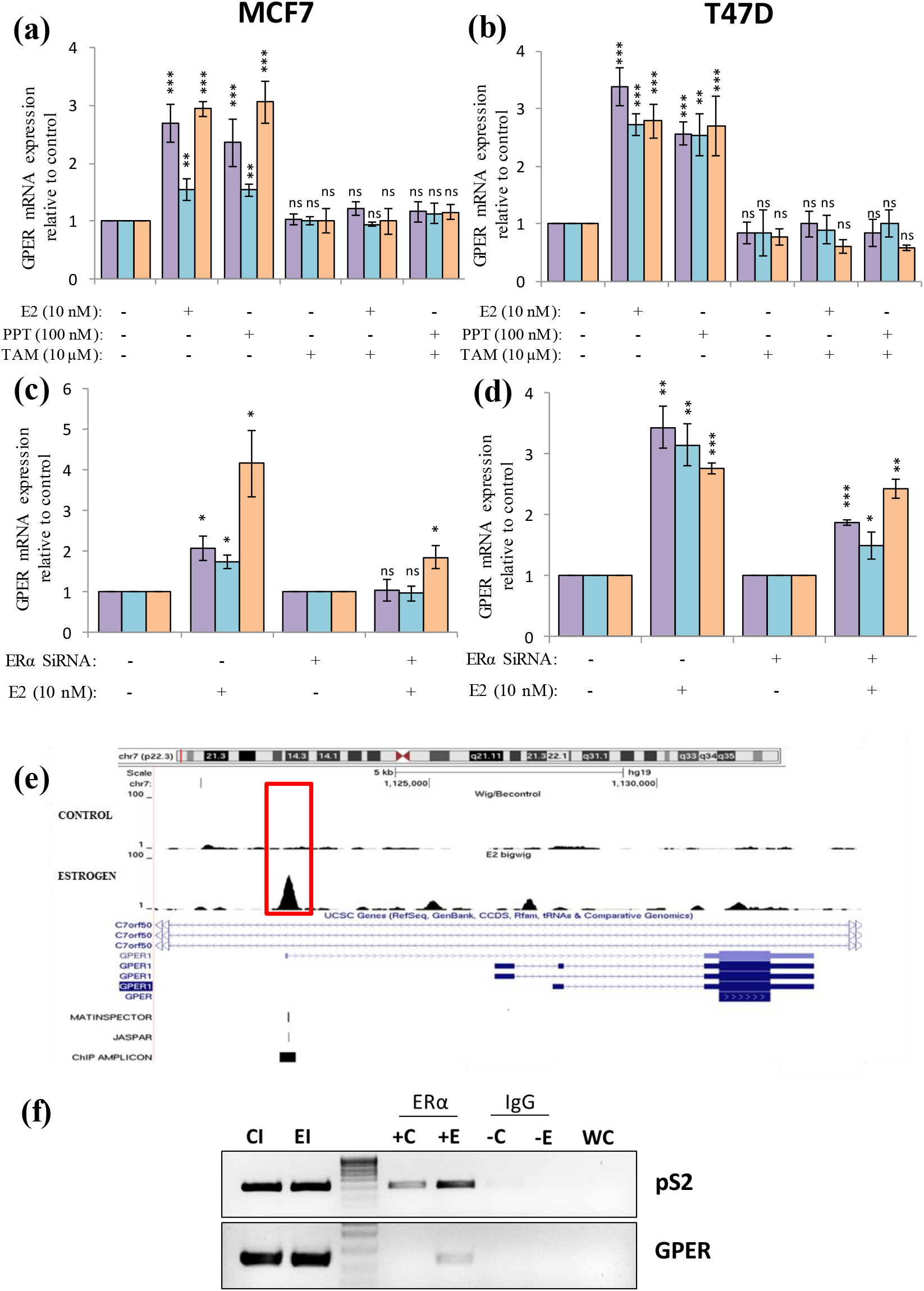
ERα-dependent induction of GPER mRNA by E2 in breast cancer cells. (a, b) Tamoxifen blocks E2-or PPT-induced expression of GPER mRNA variants. MCF-7 (a) or T47D (b) cells were stimulated with 10 nM E2 or 100 nM PPT, alone or in combination with 10 μM tamoxifen. Relative GPER mRNA variant expression levels were ascertained by RT-qPCR as in dose-response experiments. Cells treated with tamoxifen alone were also included in the experiment design. Expression levels in vehicle controls (ethanol treated cells) were set to 1 and those in E2 treated groups were expressed relative to control. Bars represent mean ± sd (n = 3). Data were subjected to ANOVA with Tukey’s HSD. *p < 0.05, **p < 0.01, ***p < 0.001. (c, d) ERα-knockdown blocks E2-mediated induction of GPER mRNA variants. MCF-7 (c), or T47D (d) cells, transfected with scrambled or ERα-specific siRNA for 24 h were stimulated with vehicle or 10 nM E2 for 72 h. GPER mRNA variant expression relative to RPL35a was ascertained by RT-qPCR. For scrambled, or ERα-specific siRNA transfected groups, the relative GPER expression levels in E2 treated cells were expressed relative to vehicle control (ethanol-treated), and analyzed by Welch two-sample t tests. Bars represent mean ± sd (n = 3). *p < 0.05, **p < 0.01, ***p < 0.001. (e) ERα binding sites at the GPER locus in the human genome. (e) ChIP-seq data obtained from MCF-7 cells treated with vehicle, or 10 nM E2 (SRA accession no. ERP000380) was analysed using Galaxy. ERα enriched regions were visualized in the UCSC Genome Browser. Note the major peak of ERα binding indicated inside the red rectangle, observed only in estrogen treated sample. An ERE was also predicted within this peak region when analysed using MATINSPECTOR or JASPAR, as indicated in the respective tracks. (f) Validation of ERα binding using ChIP assay. Sonicated chromatin samples from vehicle or 10 nM E2-treated MCF-7 cells were immunoprecipitated with non-specific IgG or ERα-specific antibody. Immunoprecipitated DNA was subjected to PCR using primer pairs designed to specifically amplify the region indicated by black bar (ChIP amplicon track). Colored bars correspond to GPER-v2 (purple), GPER-v3 (light blue), and GPER-v4 (orange) variant mRNAs.

## Discussion

Genes regulated by ERα, or those that constitute ERα co-expression network are key to understanding phenotypic differences between ER-positive, and ER-negative tumors. On this proposition, the differential screening of MCF-7 and MDA-MB-231 cDNA libraries by Carmeci et al. was predicated, which led to the identification of GPER as an ERα co-expressed marker in breast cancer cell lines and tumors [4]. Although the co-expression is confirmed by others [27], the clinical data on the association between ERα and GPER expression are inconsistent. Positive [16, 17, 21], negative [19, 22], and no association [15, 23, 24] were reported in independent cohorts. Here, using an assortment of methods to determine, and analyze GPER mRNA and protein expression in breast tumors from three independent cohorts, we found a positive association.

The TCGA-BRCA breast tumors showed bimodal distribution of ERα. By modeling breast tumors as a mixture of two Gaussian populations, it was possible to classify the tumors into ERα-low, and ERα-high groups. ERα expression varies with patient age or menopausal status, and ERα-positive tumors are more frequent in postmenopausal women [39]. Difference in age or menopausal status, arguably, could explain the existence of ERα-low and ERα-high subgroups. Consistent to this, a significantly lower ERα expression was observed in breast tumors from young (less than median age) versus old (greater than median age), or premenopausal versus postmenopausal women (supplementary data 5). On the contrary GPER mRNA expression did not differ; inconsistent with the positive correlation between ERα and GPER. However, this can be explained by the estrogen-mediated induction of GPER expression, as shown in this study. Despite low levels of ERα, the higher circulating levels of estrogen in young or premenopausal patients may raise the expression of GPER, thereby explaining similar levels of GPER in older and young, or premenopausal and postmenopausal patients.

The molecular basis of GPER-ERα co-expression has remained elusive. Carmeci et al. have discussed the possible role of AP1 and AP2 transcription factors [4]. In response to EGF and TGFα, the expression of GPER is upregulated in Ishikawa and TAM-R MCF-7 cell lines via the EGFR/MAPK signaling pathway, which involves recruitment of c-fos to the AP1 site of the GPER locus [40]. Demarco et al. reported that the IGF-IR/PKCd/ERK/c-fos/AP1 transduction pathway is involved in the IGF-mediated stimulation of GPER expression [41]. GPER promoter also contains an AP2 binding site [4]. AP1 and AP2 are also involved in the expression of ESR1, the gene encoding ERα [42, 43]. Thus, diverse signalling inputs likely coordinate GPER and ERα expression via activation of common transcription factors that regulate their expression. GPER expression in about 50% of ERα-negative breast tumors [24] implies that co-ordinated expression may not be the only basis for co-expression. Induction of GPER expression by ligand-activated ERα, as shown here, presents an alternative basis for GPER-ERα co-expression in ERα-positive breast cancer cells. Even in the absence of the natural ligand, overt growth factor signalling leading to ligand independent activation of ERα [44] may also induce GPER, thereby explaining their co-expression.

Despite volumes of clinical and cell biological data, little is known about estrogen-mediated transcriptional regulation of GPER, let alone the contribution of ERα in its expression and function. Reporting on the discovery of GPER cDNA (then referred to as GPCR-Br), Carmeci et al. discussed the non-effect of serum stimulation on its expression in MCF-7 breast cancer cells [4]. On the contrary, we found higher expression of GPER mRNA and protein in MCF-7 cells cultured in phenol red-containing medium supplemented with routine serum, compared to those cultured in phenol red-free medium supplemented with charcoal stripped serum (supplementary data 6); strongly favouring estrogen regulation of GPER. Carmeci et al. have discussed decrease in GPER expression post estrogen exposure in MCF-7 cells, which was contradicted by Ignatov et al. with their demonstration of enhanced GPER protein expression following estrogen treatment [13]. In SKBr3 cells, a widely accepted ERα-negative breast cancer cell line, estrogen and progesterone were shown to induce the expression of GPER mRNA and protein by Thomas et al., postulating the involvement of genomic nuclear progesterone receptor and non-genomic estrogen signalling [6]. Here, progesterone had a greater effect than estrogen. Ahola et al. showed that medroxyprogesterone acetate can upregulate GPER in MCF-7 breast cancer cells. However, their experiments were conducted in media containing 1 nM E2, obscuring the effect of estrogen [45]. Estrogenic regulation is also indicated by higher GPER expression in diestrus hamster endometrium [46]. However, our *in vivo* data shows higher GPER expression in murine mammary tissues, in the estrogen dominant (estrus) phase of the reproductive cycle. Taken together the present study demonstrates the positive influence of estrogen on GPER expression, both *in vivo* and *in vitro*.

In the context of estrogen-mediated induction, the binding of ERα to the upstream regulatory region of GPER, is a significant insight. The ChIP-seq and chromatin immunoprecipitation data show that GPER is a genomic target of estrogen-ERα signalling. This is not the first instance of transcriptional activation or involvement of ERα in the regulation of GPER expression. Hypoxia transcriptionally induces the expression of GPER via HIF1α binding to the hypoxia responsive element (HRE) in GPER [47]. Interestingly, the induction of GPER via IGF-IR/PKCd/ERK/c-fos/AP1 transduction pathway involves recruitment of phospo-ERα, to the AP1 site in the GPER promoter [41]. However, the regulatory regions operative in these instances are distinct from each other. This suggests that different signalling pathways co-opt different regulatory regions to alter GPER expression.

The knowledge of GPER expression in breast tumors has prognostic or therapeutic value. Being an estrogen receptor, its presence in ERα-negative tumors implies estrogen responsiveness. Given, that GPER activation can lead to cell proliferation, or cell cycle arrest, its potential as an independent prognostic marker or a therapeutic target in ER-negative breast cancer cannot be overemphasized. The significance of GPER-ERα co-expression, however, has not been adequately addressed, although it is appealing on several counts. Analysis of survival data revealed that high GPER expression is associated with significantly longer overall survival of patients with ERα-positive breast tumors. In contrast, patients with ERα-negative tumors with higher GPER expression, have poor overall survival. Thus, GPER-ERα co-expression entails favourable prognosis. 15-20% of ERα-positive breast tumors do not respond to endocrine therapy [48]. Thus, the ability to distinguish between endocrine responsive, and non-responsive ERα-positive breast tumors has therapeutic value. PR, a downstream target of ERα and a gene induced by estrogen-ERα signalling is a classical marker of estrogen responsiveness, and estrogen responsive breast tumors. Given that GPER is a downstream transcriptional target of estrogen-ERα signalling axis, GPER presents itself as a potential alternative marker of endocrine therapy response.

Estrogen-mediated induction of GPER has cell biological implications. GPER signaling activates the EGFR-MAPK pathway, which in turn activates the unliganded ERα, due to the phosphorylation of serine 118 [11]. This cross-talk between GPER and ERα is a proof of the functional interaction that is made possible by their co-expression. Estrogen-mediated induction of GPER via ERα shows that the cross-talk is bidirectional; each impacting the expression or function of the other. It also implies that in the face of estrogenic stimulation cells will be rendered more responsive to GPER activating ligands. Besides, it has greater cell biological implications in the context of breast cancer development and progression. Given the ambivalence in the data on effects of GPER activation, which indicate both promotion and inhibition of cell proliferation, we envisage dual role for enhanced GPER expression. In view of the negative impact of GPER activation on cell proliferation [49], estrogen induction of GPER may serve to produce a balancing effect to counteract the pro-proliferative effects of activated ERα. Such a mechanism would prevent excessive proliferation of the mammary epithelium in the face of estrogenic stimulation, thereby preventing tumorigenesis. On the other hand, in the light of pro-proliferative effects of activated GPER [50, 51], and higher expression of GPER in tamoxifen resistant cells [52], estrogen induction of GPER may render ERα dispensable for growth and proliferation of the mammary epithelium. Such a mechanism would subserve the emergence of endocrine therapy resistant breast tumors.

## Acknowledgements

The authors thank the Department of Biosciences and Bioengineering, IIT Guwahati, Department of Pathology, North Eastern Indira Gandhi Regional Institute of Health & Medical Sciences, Shillong, Molecular and Cellular Biology Laboratory, National Institute for Research in Reproductive Health, Mumbai, and Dr. Bhubaneshwar Borooah Cancer Institute, Guwahati, for infrastructural support. The authors acknowledge the support from DBT Centre for Molecular Biology and Cancer Research at BBCI, Guwahati. The part of the results published here are based upon data generated by The Cancer Genome Atlas (TCGA) Research Network: https://www.cancer.gov/tcga. Author thanks IIT-Bombay for Zeiss Confocal facility and Manalee Surve for assistance while imaging.

## Statements & Declarations

### Author Contributions

Material preparation, data collection and analysis were performed by Uttariya Pal, Mohan Manjegowda, Neha Singh, Snigdha Saikia, Betty S. Philip, Deep Jyoti Kalita, Avdhesh Kumar Rai, Anupam Sarma, The work in respective collaborating institutions were supervised by Vandana Raphael, Deepak Modi, Amal Chandra Kataki and Anil Mukund Limaye. The first draft of the manuscript was written by Mohan Manjegowda and Anil Mukund Limaye and all authors commented on previous versions of the manuscript. All authors read and approved the final manuscript. Uttariya Pal and Mohan Manjegowda are co-first authors in this manuscript.

### Data Availability

The TCGA-BRCA dataset analysed during the current study are available in the TCGA repository, https://www.cancer.gov/tcga

### Ethics approval

The Ethics Committee of Dr Bhubaneshwar Borooah Cancer Institute, Guwahati approved the clinical work Reference no. BBCI-TMC/SC/Appr/15/2019 and BBCI-TMC/Misc-119/3180/2018.The institute ethics committee of North Eastern Indira Gandhi Regional Institute of Health & Medical Sciences, Shillong approved to conduct IHC study on breast cancer tumor samples (NEIGR/IEC/2013/12). The animal study was approved by institution animal ethics committee of National Institute for Research in Reproductive and Child Health, Mumbai (IAEC Project No. 1/16).

### Consent to participate

Informed consent was obtained from all individual participants included in the study.

### Consent to publish

Informed consent was obtained from all individual participants included in the study.

