## Supplementary data 1 for "The G-protein-coupled estrogen receptor, a gene co-expressed with ERα in breast tumors, is regulated by estrogen-ERα signalling in ERα positive breast cancer cells"

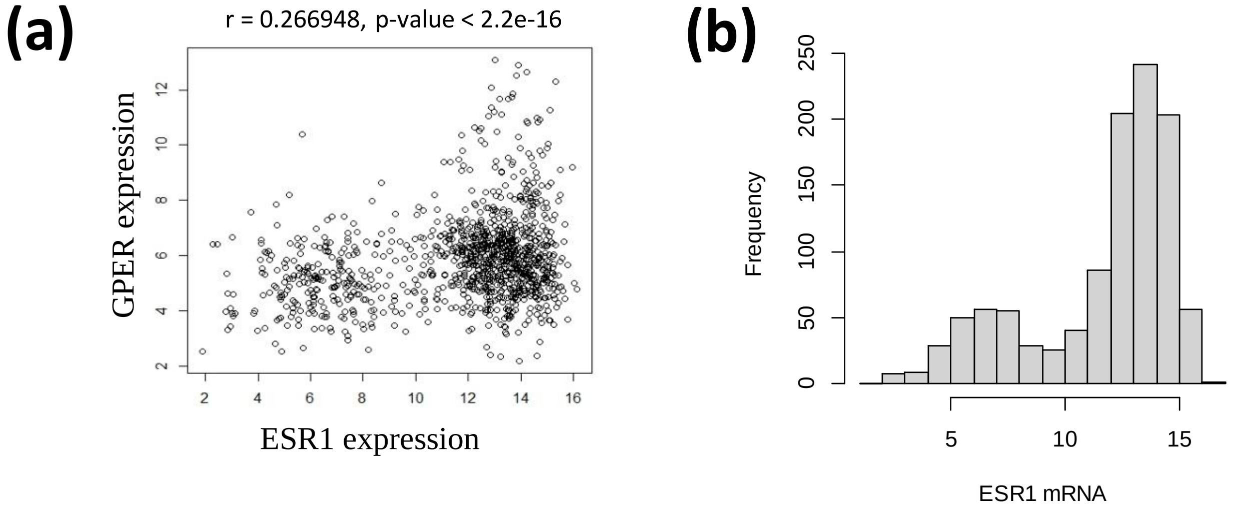


**Supplementary data 1**: (a) Correlation plot between GPER and ERα mRNA expression in primary breast tumors from the TCGA-BRCA dataset. (b) Histogram depicting the bimodal distribution of ER mRNA expression in primary breast tumors from the TCGA-BRCA dataset.
