## Supplementary data 2 for "The G-protein-coupled estrogen receptor, a gene co-expressed with ERα in breast tumors, is regulated by estrogen-ERα signalling in ERα positive breast cancer cells"

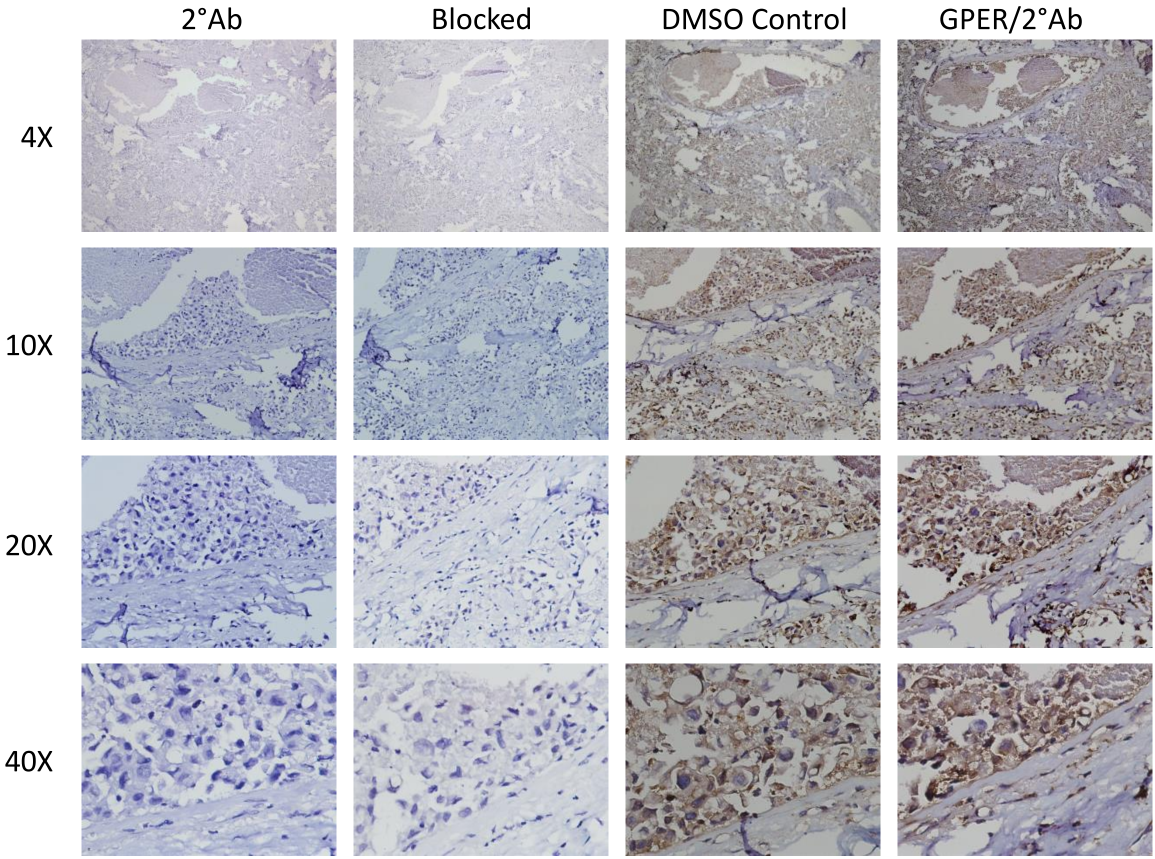


**Supplementary data 2:** No primary antibody (-ve control) or anti-GPER antibody pre-incubated with the peptide (Blocked antibody) or peptide resuspension buffer for 1 h (resuspension buffer control), or anti-GPER antibody (+ve control).
