## Supplementary data 3 for "The G-protein-coupled estrogen receptor, a gene co-expressed with ERα in breast tumors, is regulated by estrogen-ERα signalling in ERα positive breast cancer cells"

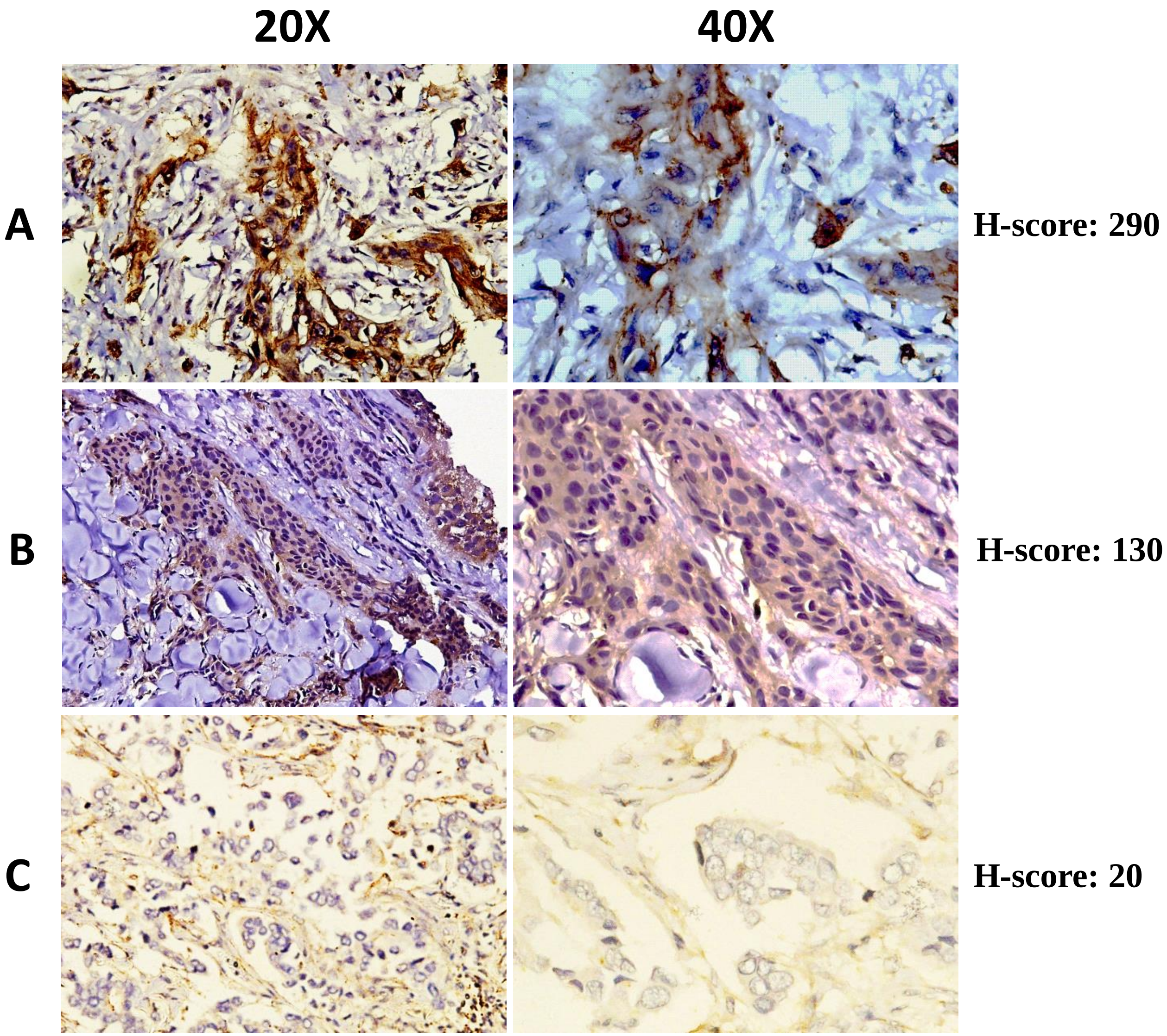


**Supplementary data 3:** Expression of GPER in the breast cancer. IHC was performed on formalin fixed, paraffin embedded breast cancer tissue sections using, anti-GPER antibody. A total of 65 cases were analyzed. Representative images for staining of high (A), medium (B) and low (C) GPER expression are shown. The corresponding H-score for high, medium, and low groups was 290, 130, and 20, respectively. Images in the first column were taken under 20× objective and those in the second column were taken under 40× objective.
