## Supplementary data 4 for "The G-protein-coupled estrogen receptor, a gene co-expressed with ERα in breast tumors, is regulated by estrogen-ERα signalling in ERα positive breast cancer cells"

**Blot - I**

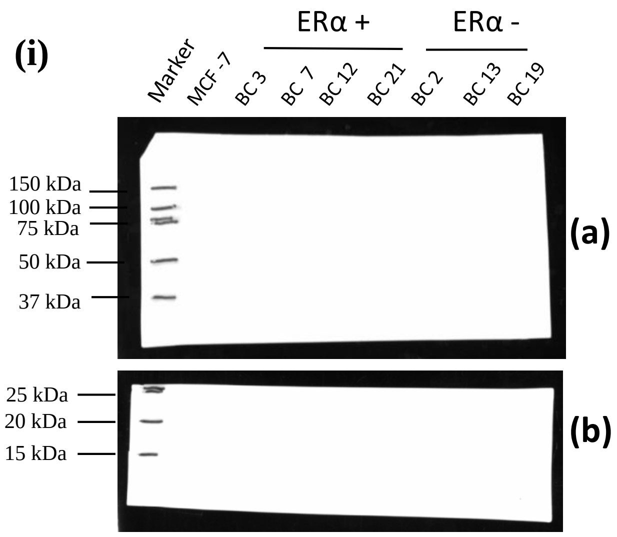


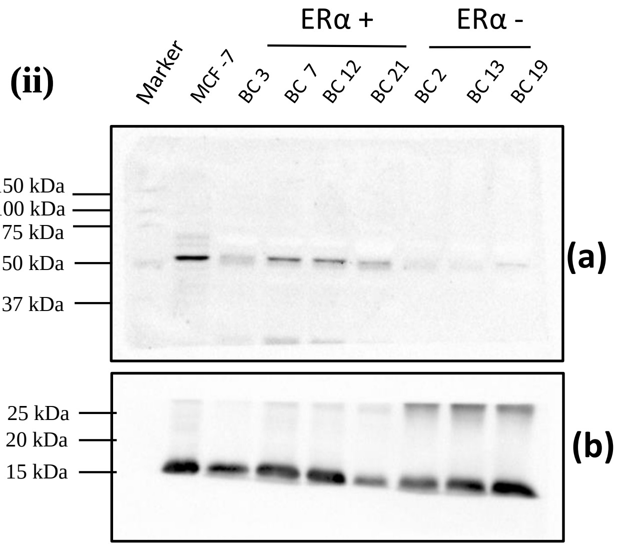


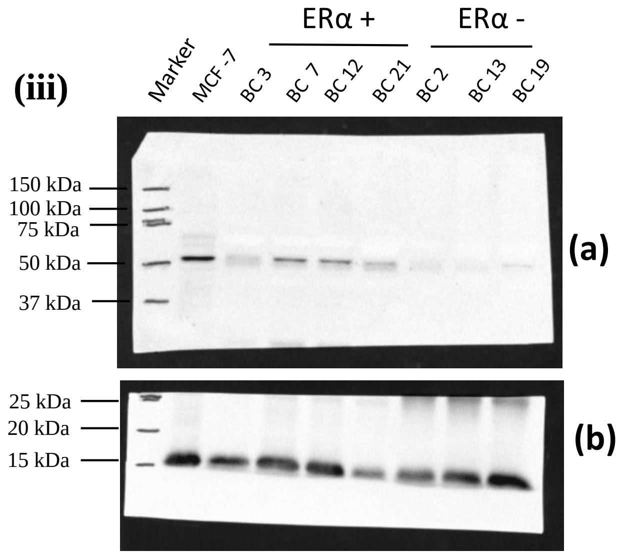


**Supplementary data 4: GPER and Histone (H3) protein expression in breast tumor samples**. Total protein was isolated from the indicated breast tumor samples using Trizol, was separated by 10 % SDS-PAGE and transferred on to nitrocellulose membrane. White light image of the blot was captured (panel i). The blot was probed for GPER (blot a) and H3 (blot b). panel (ii) shows the images of the chemiluminescence signal captured for GPER and H3, panel (iii) shows the merged images of white light and chemiluminescence.

### Blot - II

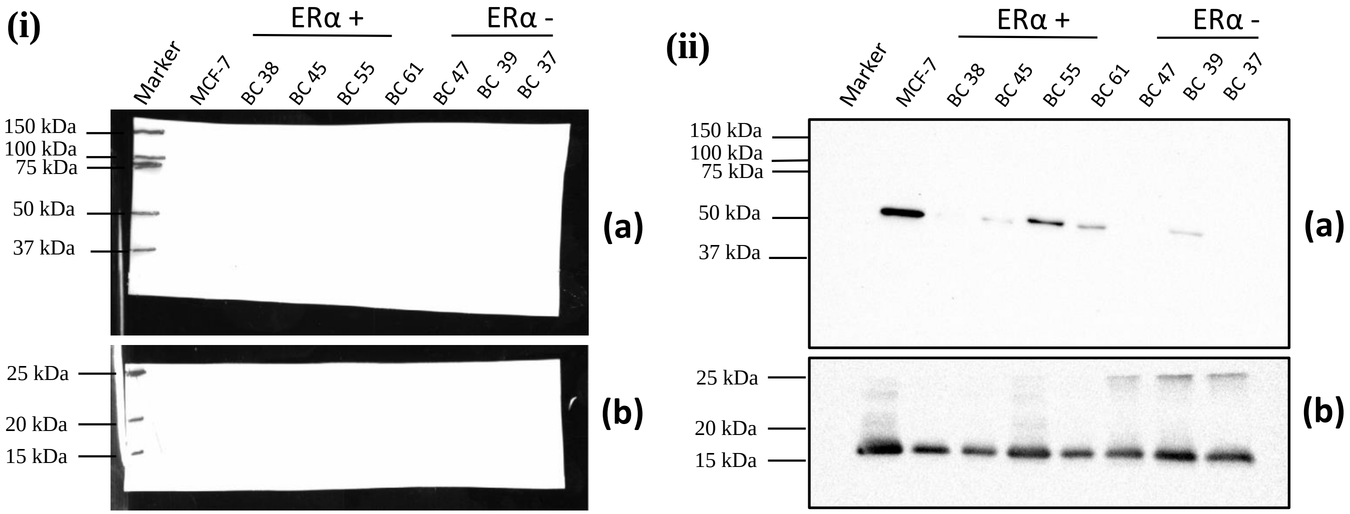


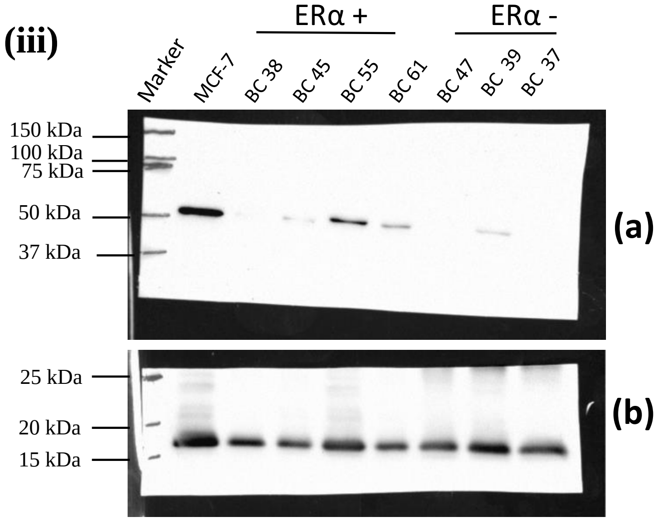


**Supplementary data 4: GPER and Histone (H3) protein expression in breast tumor samples**. Total protein was isolated from the indicated breast tumor samples using Trizol, was separated by 10 % SDS-PAGE and transferred on to nitrocellulose membrane. White light image of the blot was captured (panel i). The blot was probed for GPER (blot a) and H3 (blot b). panel (ii) shows the images of the chemiluminescence signal captured for GPER and H3, panel (iii) shows the merged images of white light and chemiluminescence.

#### Blot - III

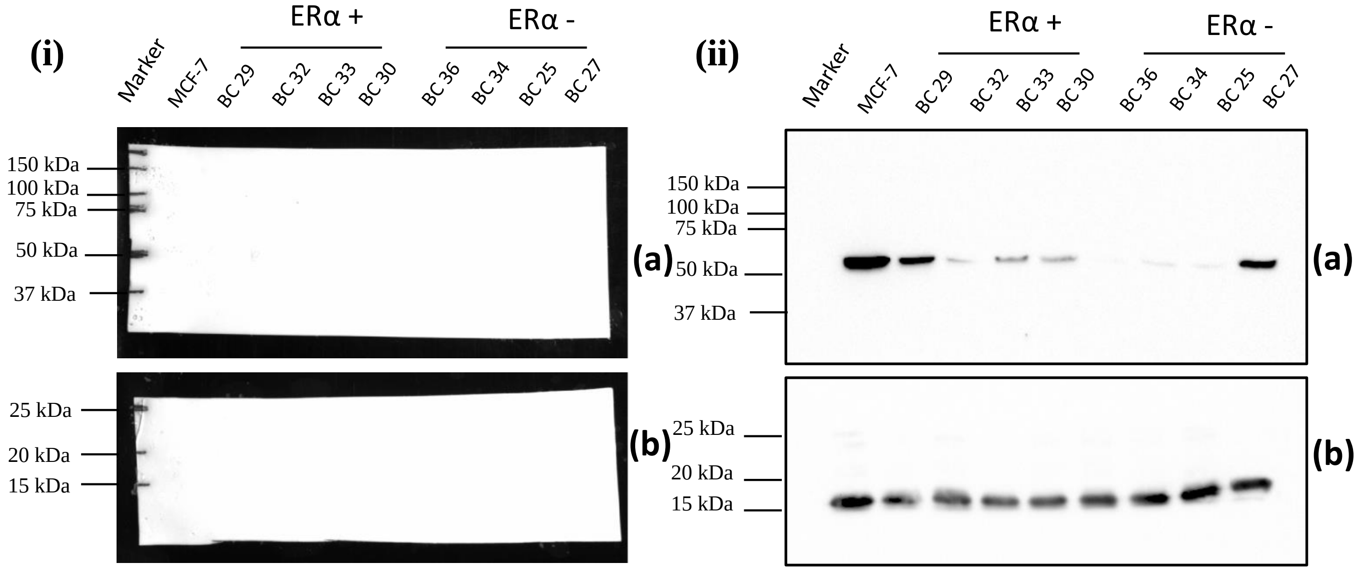


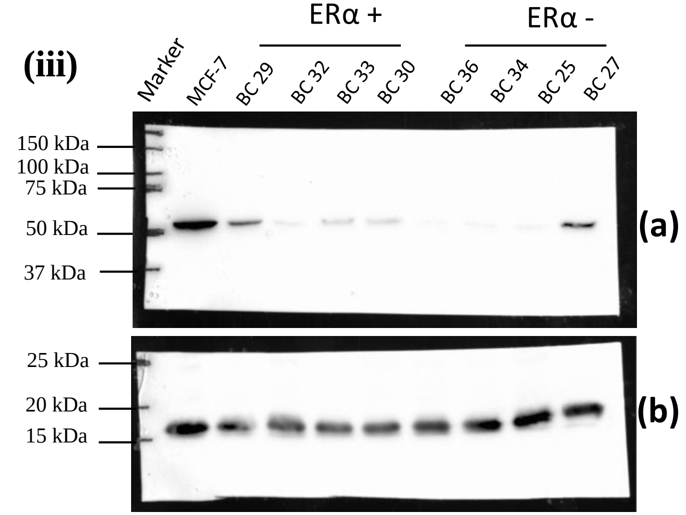


**Supplementary figure 4: GPER and Histone (H3) protein expression in breast tumor samples**. Total protein was isolated from the indicated breast tumor samples using Trizol, was separated by 10 % SDS-PAGE and transferred on to nitrocellulose membrane. White light image of the blot was captured (panel i). The blot was probed for GPER (blot a) and H3 (blot b). panel (ii) shows the images of the chemiluminescence signal captured for GPER and H3, panel (iii) shows the merged images of white light and chemiluminescence.
