## Supplementary data 5 for "The G-protein-coupled estrogen receptor, a gene co-expressed with ERα in breast tumors, is regulated by estrogen-ERα signalling in ERα positive breast cancer cells"

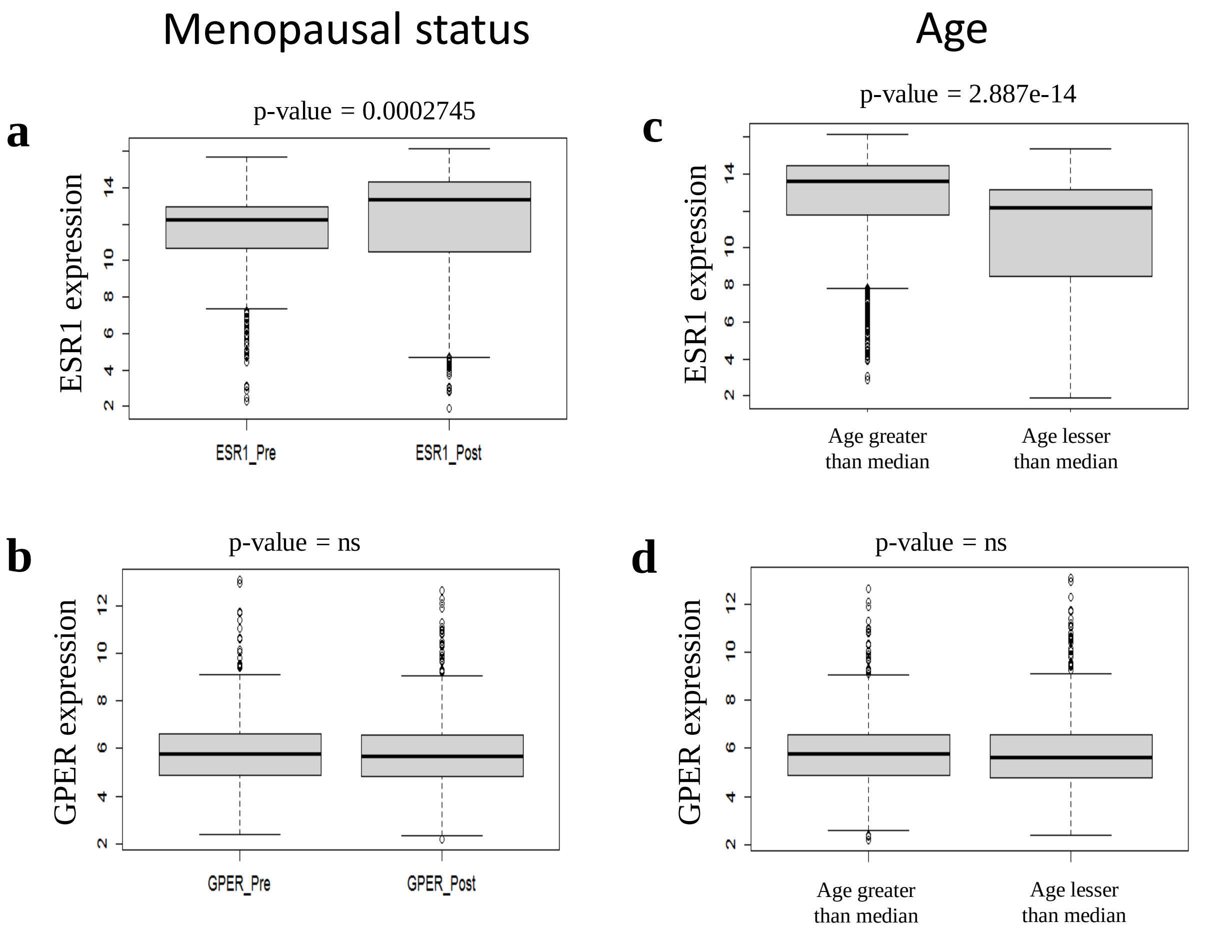


**Supplementary data 5:** Association of GPER and ESR1 with menopausal status and age in breast tumor samples (a,b) Box plots depicting the distribution of ESR1 (a) and GPER (b) mRNA expression in Post and Premenopausal breast tumor samples. (c,d) Box plots depicting the distribution of ESR1 (c) and GPER (d) mRNA expression in Age –greater than median and –lesser than median breast tumor samples. The difference in the mean GPER and ESR1 expression in two groups was analyzed by Welch two-sample t-test. p-value is mentioned above the box-plots. ns = not significant
