## Supplementary data 6 for "The G-protein-coupled estrogen receptor, a gene co-expressed with ERα in breast tumors, is regulated by estrogen-ERα signalling in ERα positive breast cancer cells"

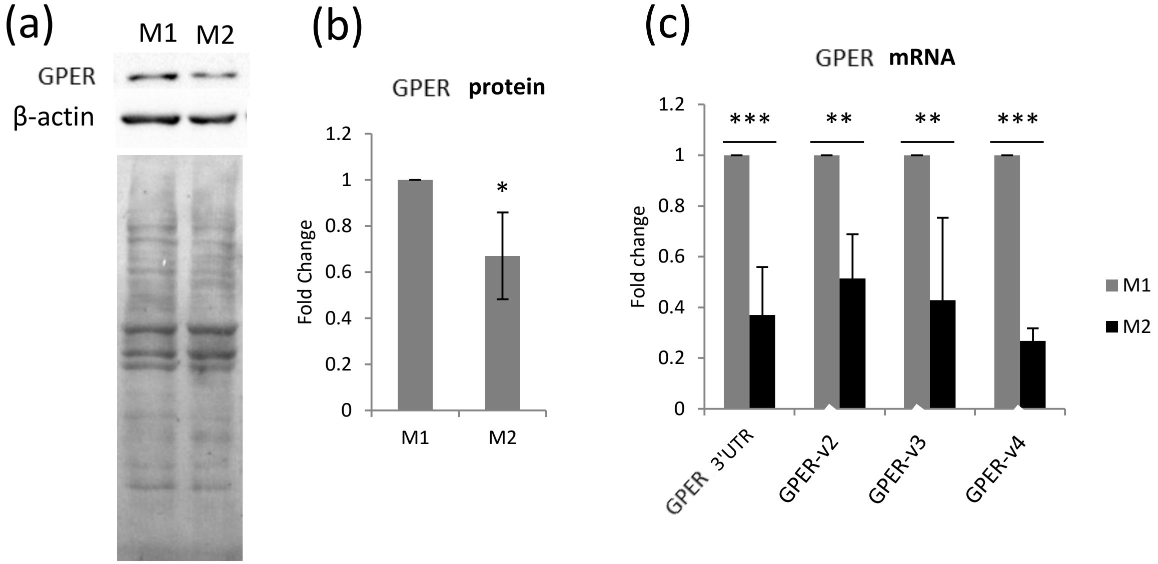


**Supplementary data 6: GPER expression in MCF-7 cells cultured in M1 or M2 media.** MCF-7 cells were seeded into 35mm dishes and allowed to grow for 48 h. Cells were then cultured in M1 or M2 medium for another 72 h, with medium replenishment after 48 h. Cells were lysed in TRIzol. 30 µg of total protein was used in western blotting to assess the GPER (a). The bands were quantified and normalized against ponceau S (b). Total-RNA was isolated and cDNA was synthesized from 2 µg of DNaseI treated total RNA. cDNA equivalent to 20ng of total-RNA was taken as template in the qRT-PCR reactions. RPL35A was used as an internal control. The relative expression levels of the GPER mRNA variants were analyzed by ΔΔCt method and expressed as fold change in M2 with respect to M1 (c). The bars in the graphs represent the mean fold-change ±SD, n=5 for A, n=6 for C, * $p< 0.05$, ** $p<0.01$, *** $p<0.001$.
